# Efficient Agony Based Transfer Learning Algorithms for Survival Forecasting

**DOI:** 10.1101/2021.02.24.432695

**Authors:** Abhinav Tamaskar, James Bannon, Bud Mishra

## Abstract

Progression modeling is a mature subfield of cancer bioinformatics, but it has yet to make a proportional clinical impact. The majority of the research in this area has focused on the development of efficient algorithms for accurately reconstructing sequences of (epi)genomic events from noisy data. We see this as the first step in a broad pipeline that will translate progression modeling to clinical utility, with the subsequent steps involving inferring prognoses and optimal therapy programs for different cancers and using similarity in progression to enhance decision making. In this paper we take some initial steps in completing this pipeline. As a theoretical contribution, we introduce a polytime-computable pairwise distance between progression models based on the graph-theoretic notion of “agony”. Focusing on a particular progression model we can then use this agony distance to cluster (dis)similarities via *multi-dimensional scaling*. We recover known biological similarities and dissimilarities. Finally, we use the agony distance to automate transfer learning experiments and show a large improvement in the ability to forecast time to death.

## 1 INTRODUCTION

Cancer progression modeling is a mature subfield of cancer informatics [7]. The desirable models seek to recapitulate or forecast the accumulation of genomic events in the course of a patient’s disease. Given these purposes, progression models often take the form of hierarchical combinational structures such as phylogenetic trees [1][14] or various forms of Bayesian networks [3, 11]. In this paper we consider a cancer progression model (CPM) to be a directed acyclic graph (DAG) defined over a collection of (epi)genomic events. This view encompasses the structures of both phylogenetic trees and Bayesian networks and is agnostic with respect to probabilistic assumptions or interpretations of any particular CPM.

Research on CPMs often focuses on accurately recreating an underlying ground truth. Most research papers involve the presentation of a new algorithm and showing empirically or rigorously that this algorithm reconstructs a latent CPM correctly. These methods are, for the most part, retrospective and are predicated on the theory that understanding the course of evolution of a particular patient population will shed insight into the nature of that particular cancer and, hopefully, its treatment.

To our knowledge CPMs have not yet made this final clinical leap. In particular while similarities between CPMs have been explored via edit distances [15], the similarity between progression models across different cancer types have not been exploited. As cancer progression seems to correlate with phenotype it follows that patients with similar disease progressions will be similar in terms of clinical presentation.

This paper contains two main contributions that address these issues. In order to compare progression models across different cancer types we *first* introduce a notion of similarity based on the graph theoretic concept of “agony.” This is an alternate notion of distance that measures the preservation of structural (involving driver genes) relationships in a progression model. Thus the semantics of looking at agony directly correspond to the orderings of events given by two CPMs.

The *second* contribution of this paper involves using this notion of distance to automate transfer learning, with specific experiments directed towards survival forecasting. In transfer learning one seeks to leverage the learned capability to perform *source task* to improve the ability to perform a *target task*. Usually the choice of source and target based on the fact that they are in some ways similar. Here we assume that similar progression models correspond to similar phenotypes. We fix the target task as forecasting survival time for a particular cancer and then choose the source task as predicting survival from the cancer which has the smallest agony based distance from the target task Figure 1. We show empirically that our clusters correspond to meaningful biological similarities and that agony-guided transfer learning significantly improves performance in some cases.

**Figure 1:**
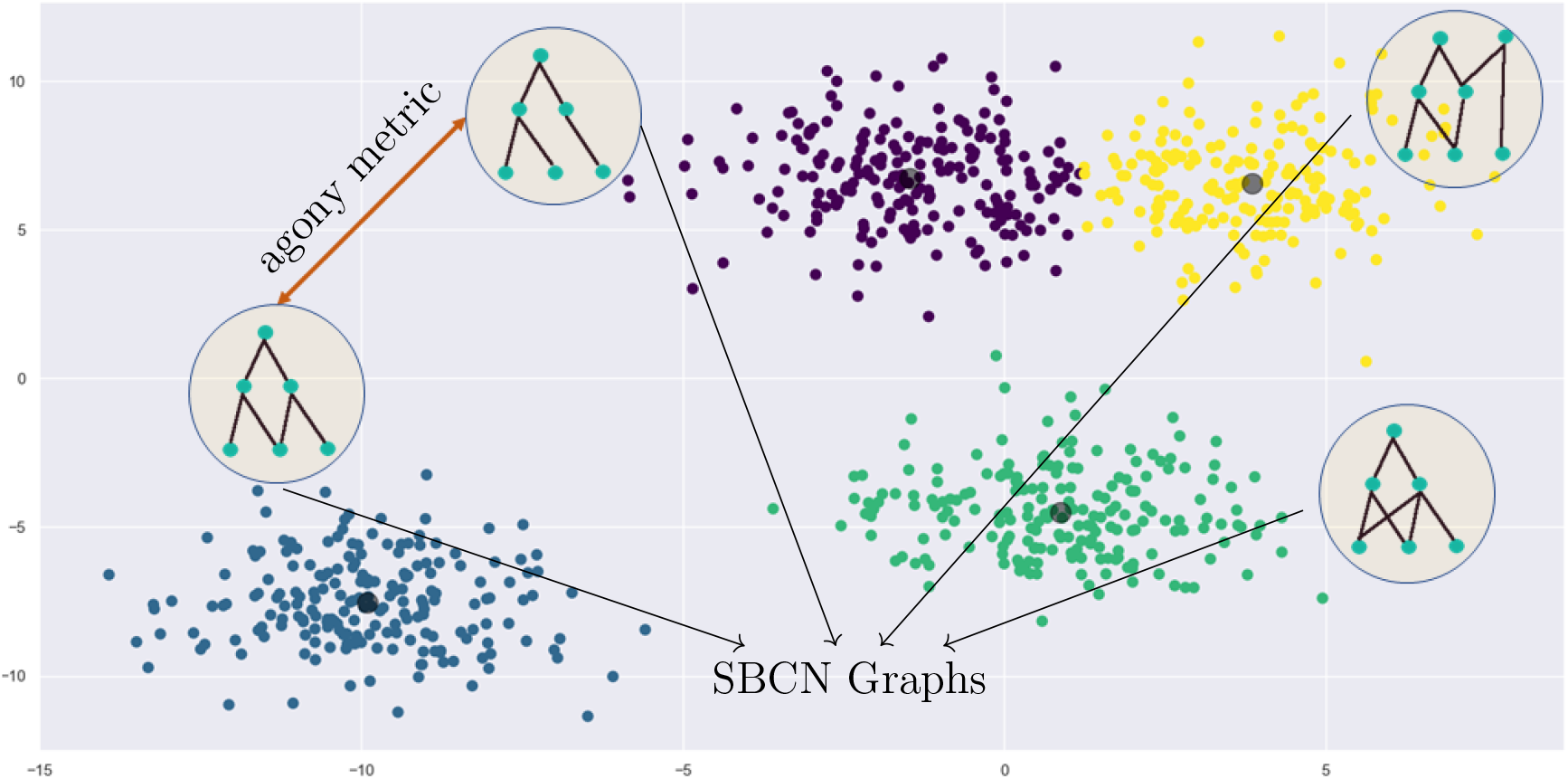
Patient clusters along with their respective SBCNs. Each cluster has its own progression model, which are then used to calculate the distance between clusters. This in turn is used to automate transfer learning.

The rest of the paper is structured as follows. In Section 2 we review the necessary background material. In particular we review progression models, survival forecasting, and transfer learning and we introduce the notion of *agony* a way of measuring the similarity of two progression models. Then in Section 3 we report on two experiments in using agony for bioinformatic purposes. The first experiment consists of clustering patients in different cancers using pairwise agony dissimilarity. The second experiment automates source task selection in transfer learning using minimum-agony distance as defined in Section 2. Finally in Section 4 we provide a discussion and pointers to future work. We include an online appendix where we include all technical details ^1^.

## 2 BACKGROUND

### 2.1 Cancer Progression Modeling

For many years, the dominant paradigm for understanding cancer has been one of continual stochastic mutation and selection. In this process some mutations — called *driver* mutations — are culpable in continued proliferation and the acquisition of phenotypic *hallmarks* [12] while others — so called *passengers* — are mere incidental alterations that are preserved via clonal expansion. In cancer genomics two problems go in to progression modeling. The first is the identification of driver mutations from the whole of genetic information. This has been addressed by efforts such as The Cancer Genome Atlas and the Catalogue Of Somatic Mutations In Cancer (COSMIC). The second task, and the one with which we concern ourselves, is the task of reconstructing the history in which driver mutations were acquired.

For the purposes of this paper we focus on a particular progression model where the partial order is due to a particular *selective advantage relation*. This relationship is called *prima facie* causality because it was originally developed as such in the philosophical literature by Patrick Suppes [24]. For this reason we keep the term causality for our progression models but remark that it is the close correspondence between this relation and the evolutionary accumulation of mutations that is of interest to us.

#### Definition 1

(Prima-Facie Causality). An event ***c*** is a *prima facie cause* of another event ***e*** if the following two conditions hold:

- (Temporal Priority) ***t_c_** < **t_e_***, where ***t***_*_ is the time of observation of an event.
- (Probability Raising) **Pr[*e*|*c*]** > **r[*e*|¬-*c*]**.

Two remarks are in order. The first is that the notion of prima facie causality here is different from the type of causality more commonly considered in the computer science literature [21] using counter factual possible worlds or do-calculus or structural equations.

However, it is not entirely distinct and the probability raising condition maps to the same counterfactual intuitions that underlie clinical trials and the “do calculus.” An additional and critical similarity is that we are able to construct causal graphs, a topic to which we return shortly.

The second remark is that this approach to causality mirrors the biological processes of the accumulation of driver genes [23]. Suppose at some time ***t*_1_** a patient undergoes a mutation in *KRAS* causing the tumor to grow rapidly. As this process continues the cells in the mass will begin to experience *hypoxia*, which necessitates an alteration of behavior to angiogenesis or to metastasis. In this condition a mutation in, for example, *VEGF*, at time ***t***_2_ > ***t*_1_** would lead to necessary metabolic changes (e.g. angiogenesis) to ensure continued growth. Observe also that without the initial mutation in *KRAS* the *VEGF* mutation would not have provided any advantage so it is presumably less likely to occur.^2^

#### Definition 2

(Suppes Bayes Causal Network). A Suppes Bayes Causal Network (SBCN) is a DAG where for every edge ***v_i_*** → ***v_j_***, Suppes conditions for *prima facie causation* hold, that is:

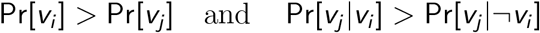

For our purposes the input to learning an SBCN is cross-sectional patient data, that is a binary matrix 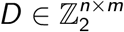 where ***n*** is the number of patients and ***m*** is the number of genes that are being considered. The learning of an SBCN structure from data can be implemented efficiently using open source software [6]. For the purposes of this paper we consider only point mutations do not account for the variant type (e.g. missense, nonsense).

While we have chosen to work with this particular notion of causality, our methods in this paper are agnostic as to the semantics of the underlying progression model. We chose Suppes causation because it has previously been applied in biology and is computationally tractable for large data sets [22].

### 2.2 Graph Agony

*Agony* is one of many measures assessing hierarchies in directed graphs. Some of these are known to be computationally intractable [8]. Agony, however, is computable in polynomial time. In particular, agony is a measure of the degree to which a directed graph is acyclic [25, 26]. To our knowledge it is the only computationally tractable method for comparing pairs of directed acyclic graphs, and yet approximates as close as other graph distance functions.

#### Definition 3

(Rank function). A rank function on a graph *G* = (*V, E*) is a map *r*: *V* → {1…, *n*}.

Intuitively the rank function can be thought of as specifying an ordering on the nodes of the graph. In the context of a progression model the rank function could be thought of as a hypothesized temporal ordering or as assigning certain mutations to *levels* in a hierarchy.

#### Definition 4

(Agony of a Graph With Respect to a Rank Function). For a graph *G* = (*V, E*), *V* = {*v*_1_,…, *v_n_*} and a rank function *r*: *V* → {1…, *n*} the agony of *G*, with respect to *r*, is defined as

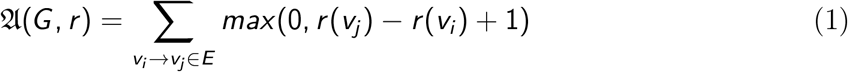

Clearly from Equation (1) the larger the difference in rank between a parent node and its child, the larger the agony. In practice a given rank function ***r*** is not available *a priori* which is an issue because the value of 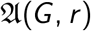 is highly dependent on ***r***. In light of this, we define the general agony of a graph as follows:

#### Definition 5

(Agony). For a graph ***G*** we define the *agony of the graph **G*** to be

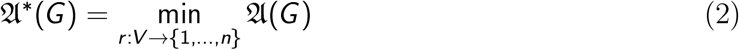

For the rest of this paper whenever we refer to the *agony of a graph* we mean it as given in Definition 5 unless otherwise stated. It is not obvious from the form of Equation (2) that 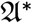 is computationally tractable. However, if one constructs the dual problem to Equation (2) one arrives at the maximal Eulerian subgraph problem, which can be solved efficiently. We refer the reader to [25] for details.

#### Definition 6

(Agony Between Graphs). For two graphs ***G_1_***, ***G_2_*** on the same set of nodes, ***V***, let ***G’*** be the union of the two graphs, without duplicate edges. The agony between ***G_1_***, ***G_2_*** is defined as

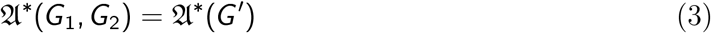

In the case where ***G_1_***, ***G_2_*** are progression models then Equation (3) can be thought of as a measure of mutual inconsistency. If there are conflicting or contradicting paths in ***G_1_*** and ***G_2_*** then 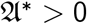Figure 2. For the case where ***G_1_***, ***G_2_*** are in fact progression models, Equation (3) possesses properties useful for comparing them.

**Figure 2:**
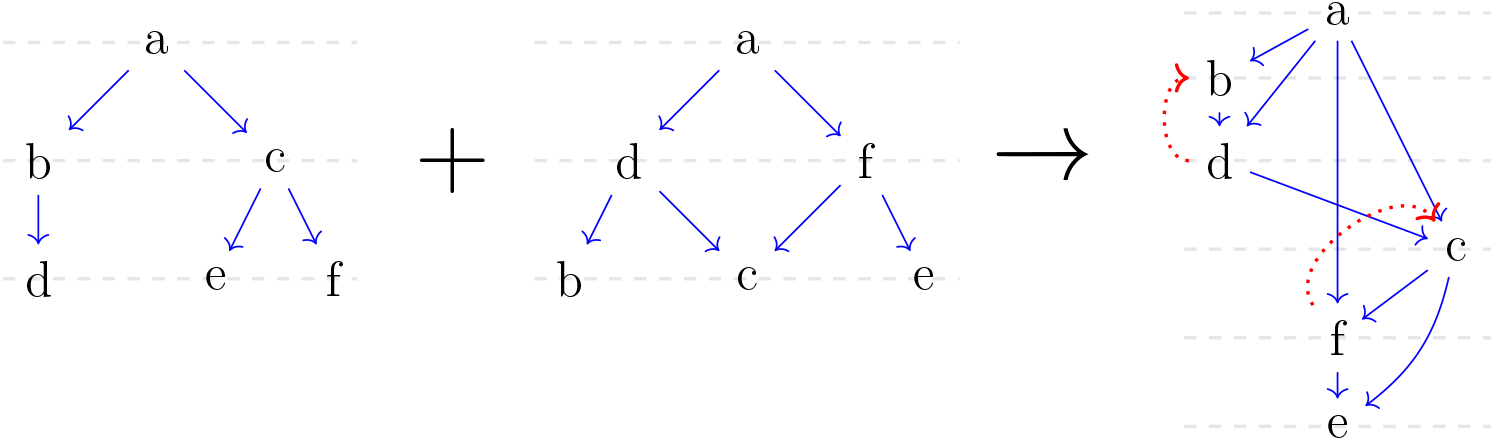
*Agony* metric between progression models. The grey lines represent the minimal rank function, and the red arrows are the edges contributing to the *agony*. In this case the agony distance is **4**.

### 2.3 Survival Forecasting

A common problem in clinical oncology — and clinical care in general — is forecasting the time until a meaningful change in the patient’s condition occurs. For example the time until death, the time until disease progression, or the time until the acquisition of a particular hallmark. Data of this type consists of a duration, which is usually measured from the beginning of an observation period (e.g. a clinical trial) until the event of interest occurs. In general a time-to-event data set 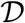 for a survival forecasting problem involves observations of the form

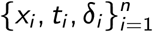

where *x_i_* = (*x_i_*_1_,…, *x_i_*_m_) is a vector of covariates, *t_i_* is the time of the event or censorship, and *δ_i_* is an indicator variable marking that the event was truly observed (***δ_i_* = 1**) or if censorship has occurred (***δ_i_* = 0**). There are many approaches to survival analysis - most either involve learning a function ***f (x)*** that returns a predictive survival time or compute a hazard ratio for a specific value of ***x***. The most well-known approach is Cox’ proportional hazard model [5] which assumes that the probability that the event occurs at time ***t*** follows an exponential distribution. We refer the reader to the two monographs [17, 19] for a thorough treatment.

The machine learning literature has also attempted to address the survival problem with standard methods such as the support vector machine [28, 10] and deep neural networks [29, 29]. For our experiments we used a deep neural network model called SurvivalNet as our survival forecasting method. We chose this method because the software had preimplemented several important neural network techniques such as drop-out and Bayesian optimization for model hyperparameters[29]^3^.

The most common method for evaluation of survival forecasting is the *concordance measure* [13] which compares the relative risk assigned to patients by the model to the order in which they actually died. Correctly ordered pairs are rewarded and incorrectly ordered pairs are penalized. We evaluate our survival forecasting models by concordance on a test set and prove the translational utility of the system.

### 2.4 Transfer Learning

Transfer learning [20, 4] is a technique in machine learning where information from a *source task* is used to improve performance on a *target task*. For example if the task is to recognize cars in natural images a transfer learning approach might be to train a system to recognize trucks and then apply it to images looking for cars. Clearly part of the consideration here is that trucks and cars are, in some ways, similar. While research in transfer learning focuses on many different aspect of the *process* of transferring knowledge [20], the source and target tasks are usually treated as given. For clinical oncology, source and target tasks should be similar in a way that is *biologically significant*. For example, in [29], the authors augment the training data for BRCA cancers with patient data from OV and UCEC because they are all hormone-driven tumors.

We generalize this by performing experiments where we augment the data by choosing the cancer with the closest (least agony) progression model. Specifically we begin with a *target task* 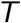 which is the survival forecasting problem for a particular cancer 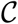, and then choose as the source task the data from the cancer 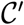 which has the smallest agony distance from 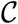.

## 3 EXPERIMENTS AND RESULTS

Our aim is to be able to evaluate the utility of agony as a (dis)similarity measure between different cancer (sub)types. To evaluate how well agony is capturing biological information we first performed two clustering experiments, detailed in Section 3.1. In order to make our approach *translational* we performed transfer learning experiments between low agony and high agony cancers, and show empirically that low agony transfer learning improves performance more than high agony. We report on these results in Section 3.2.

### Data Sources and Preprocessing

For our experiments we used the data from the TCGA 2018 PanCancer Atlas accessed via cBioPortal. This data consists of approximately 11,000 spread across 33 different tumor types. Before all experiments the mutation data from each cancer was processed into a binary matrix 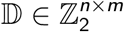 where each row corresponded to one of the ***n*** patients. The ***m*** genes were filtered to be only those that were either included in both tiers of COSMIC or those and those considered to be driver genes by TCGA.

### 3.1 Agony Recovers Biologically Meaningful Dissimilarity

#### 3.1.1 *Agony* across clusters shows biologically significant similarities

The first experiment was to explore the similarities between cancers based on their respective progression models. The first step involved fitting, for each cancer type 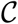, a SBCN 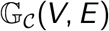. For each cancer type the top 100 most frequently mutated genes were selected and a SBCN was constructed using point mutations in these genes as the nodes. Each SBCN had approximately 1,000 edges. For each pair of graphs we calculated the *agony* distance between the pair in accordance with Equation (3). In Figure 3 we plot the pairwise agony values for each cancer type on a logarithmic scale, and show a clustering of the data using a multi-dimensional scaling (MDS) embedding of the data into two dimensional space.

**Figure 3:**
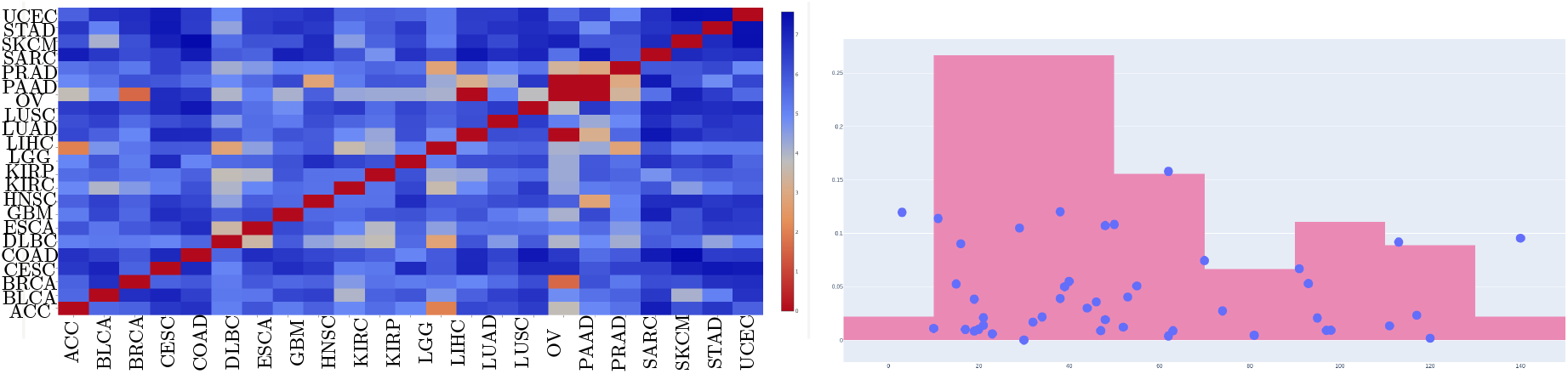
(a) Heatmap for agony distance between different cancer types. (b) Increase in concordance as a measure of the agony metric, we notice a higher concentration of pairs with an increase in concordance for low agony distance, validating our hypothesis.

We observe that skin cutaneous melanoma has a high dissimilarity to other cancers, which may be hypothesized to be due to the central role played by *BRAF* in controlling growth and apoptosis. Further analysis would be necessary to validate this hypothesis. We also note that in the MDS clustering the hormone-driven cancers UV, BRCA, and UCEC are farther from the other subtypes, which suggests the agony distance has recovered this distinction. However we expected these three to be close to each other in agony distance, which is not the case. Our hypothesis is that since BRCA has well delineated subtypes [16], due to Simpsons’ paradox [27] the overall BRCA population is not representative and that combining them loses information. We check this in the next subsection.

#### 3.1.2 *Agony* across BRCA subtypes captures known dissimilarities

To see if agony could capture well-known subtypes, and to assess if Simpson’s paradox played a roll in the overall disimilarity of BRCA to other cancers, we stratified the TCGA PanCancer BRCA patients into the five subtypes given in the data: Luminal A (LumA, ***n* = 499**), Luminal B (LumB, ***n* = 197**, Her2-Enriched (Her2, ***n* = 78**), Basal-like (Basal, ***n* = 171**), and Normal/Untyped (Normal, ***n* = 36**).

Figure 4 shows the pairwise agony distances across the BRCA subtypes. From the data in Figure 3 and Figure 4 we hypothesize that agony is recovering meaningful discrepancies in progression models and as well as meaningful *phenotypic differences*. This hypothesis motivated our use of agony as a metric for selecting the source task in transfer learning.

**Figure 4:**
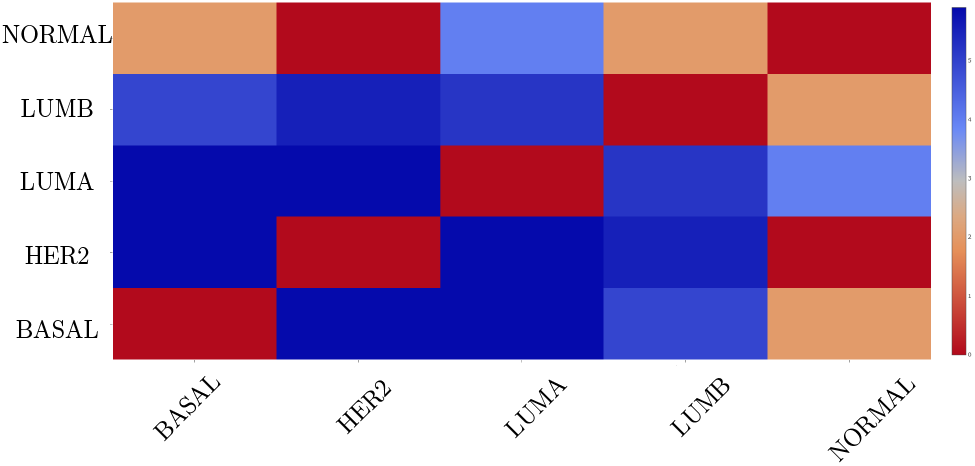
Heatmap for agony distances between the BRCA subtypes.

### 3.2 Transfer Learning

In the following experiments we split the data into train (**60%**), validation (**20%**), and test (**20%**) sets. Survival net was run with default values including automatic procedures for regularization and hyperparameter fitting. In our transfer learning experiments we did the actual transferring by adding the data from the source cancer to the training data.

#### 3.2.1 Low *agony* identifies similar cancer types, by improving concordance

We refer to fig. 3(b) to infer that most of the high concordance increase happens in the region of low agony metric, giving a confidence boost to the fact that lower agony distance is a good indicator of the accuracy gained from transfer learning.

For showcasing actual cancer pairs we chose LUAD as the dataset to be augmented. For LUAD two cancers which have a low agony distance are PRAD and MESO. These have 566, 494, and 87 patients, respectively. We performed two transfer learning experiments on these three cancers. First we evaluated SurvivalNet on LUAD alone, achieving a mean concordance 0.552 and a median concordance of 0.554 over 93 rounds of training and testing. We then augmented the LUAD training data with the data from PRAD (100 rounds), which lead to an increase in mean and median concordance to **0.735** and **0.7324** respectively. The difference in distribution means was statistically significant (Wilcoxon *p* =2e-16). To check that this was not simply the result of an increased amount of training data, we performed the same experiment with MESO (83 rounds) as the source. This analysis shows a modest increase in mean (**0.631**) and median (**0.639**) (*p* = **6.025**e-16). A figure demonstrating the improvement is given in the appendix.

#### 3.2.2 High *agony* identifies vastly different types does not improve concordance

It is possible that simply combining *any* two datasets might yield increased concordance. To test this we performed the same experiments as above but with a *high agony* pair. Specifically we performed survival forecasting on STAD (*n* = **440**) and PAAD (*n* = **184**) and then a transfer-learning with PAAD as the source. In this context survival net performed well on STAD (54 rounds) and PAAD (60 rounds) individually but in the transfer learning context (44 rounds) performance the concordance did not meaningfully change. A Wilcoxon test on the concordance of STAD and STAD+PAAD has a p-value of 0.7659, so we fail to reject the null hypothesis that the distributions are the same. We show the lack of improvement citing the results in Figure 3(b) and Table 1 for STAD+PAAD. The regression curves show a distinct decrease in concordance gain with the increase in the agony metric, in addition to which the showcased example, STAD+PAAD, shows a statistically insignificant change in the accuracy values.

**Table 1:** We report the mean and median concordances for our experiments along with bootstrap **95%** confidence intervals. A single cancer name represents running survival net on that data alone. Other rows take the form of TARGET+SOURCE. For transfer learning experiments we give the pairwise log agony distance

| Data | Mean Concordance | Median Concordance | Log Agony |
| --- | --- | --- | --- |
| LUAD | 0.5522, [0.5421, 0.5620] | 0.5542, [0.5424, 0.5663] | — |
| LUAD+PRAD | 0.7353, [0.7263, 0.7440] | 0.7324, [0.7225, 0.7389] | 3.16 |
| LUAD+MESO | 0.6317, [0.6204, 0.6432] | 0.6395, [0.6524, 0.6251] | 0 |
| STAD | 0.5714, [0.5574, 0.5855] | 0.5759, [0.5680, 0.5926] | — |
| PAAD | 0.6120, [0.5966, 0.6278] | 0.6239, [0.6524, 0.6066] | — |
| STAD+PAAD | 0.5748, [0.5672, 0.5826] | 0.5780, [0.5671, 0.5920] | 7.11891 |

While these results are not definitive, they suggest that agony is capturing a meaningful distinction *phenotypically* and could be used to guide further transfer learning experiments.

## 4 DISCUSSION

We have proposed agony as a novel method of quantifying the (dis)similarity between progression models by discovering conflicts in their hierarchical relationships. We have shown empirically that this measure recovers known biological similarities and differences in cancer types. Finally we showed the potential for clinical utility by using agony to automate the choice of a source task in transfer learning experiments. To our knowledge this is the first biological attempt to automatically solve the source-selection problem, which is of research interest in the artificial intelligence community. Our experiments showed a correspondence between low agony distance and increased task performance.

Our approach can be easily generalized. Agony clustering is agnostic to the semantics of the underlying CPM but, since it measures pairwise inconsistency, it directly accounts for the semantics of whichever CPM is chosen. Also, our transfer learning methodology is amenable to any machine learning technique. An obvious next step is to use agony to compare different progression models. Another option is to vary the machine learning task in question. One can even generalize beyond machine learning to investigate whether populations with similar progression models are similar in *any* interesting phenotypic characteristic.

In the near term we hope to expand on the theoretical foundations of agony to large graph limits and to graphons [18]. This would allow us to bring large sample theory to bear on the techniques presented in this paper, and potentially allow us to generalize SBCNs to cases of continuous variables, e.g. gene expression. Graphons have also received growing attention from the machine learning literature [2, 9]. We believe that even its current form graph agony can be a valuable tool for both clinical and research cancer bioinformaticians.

## 5 Acknowledgements

This work was supported by National Science Foundation Grants CCF-0836649 and CCF-0926166,and a 684 National Cancer Institute Physical Sciences-Oncology Center Grant U54CA193313-01 (to B.M.).

## Footnotes

https://github.com/epsilon-0/pan-cancer

We acknowledge that rigorously stating this would require specifying base-rate mutations in *VEGF* the tumor. The thrust of the argument is that there is selective pressure on the tumor as a whole to survive hypoxia and as such the clones that survive and are biopsied are those which have such an advantageous mutation.

Unlike the blackbox models from SVM and DNN, SBCN can provide an evolutionary/causal explanation for the cancer prognostics.

## References

[1] N. Aguse, Y. Qi, and M. El-Kebir. Summarizing the solution space in tumor phylogeny inference by multiple consensus trees. Bioinformatics, 35(14):i408–i416, 07 2019.

[2] E. M. Airoldi, T. B. Costa, and S. H. Chan. Stochastic blockmodel approximation of a graphon: Theory and consistent estimation. In Advances in Neural Information Processing Systems, pages 692–700, 2013.

[3] N. Beerenwinkel, N. Eriksson, B. Sturmfels, et al. Conjunctive bayesian networks. Bernoulli, 13(4):893–909, 2007.

[4] D. C. Ciresan, U. Meier, and J. Schmidhuber. Transfer learning for latin and chinese characters with deep neural networks. In The 2012 International Joint Conference on Neural Networks (IJCNN), pages 1–6. IEEE, 2012.

[5] D. R. Cox. Regression models and life-tables. Journal of the Royal Statistical Society: Series B (Methodological), 34(2):187–202, 1972.

[6] L. De Sano, G. Caravagna, D. Ramazzotti, A. Graudenzi, G. Mauri, B. Mishra, and M. Antoniotti. Tronco: an r package for the inference of cancer progression models from heterogeneous genomic data. Bioinformatics, 32(12):1911–1913, 2016.

[7] R. Diaz-Uriarte and C. Vasallo. Every which way? on predicting tumor evolution using cancer progression models. BioRxiv, page 371039, 2019.

[8] I. Dinur and S. Safra. On the hardness of approximating minimum vertex cover. Annals of mathematics, pages 439–485, 2005.

[9] J. Eldridge, M. Belkin, and Y. Wang. Graphons, mergeons, and so on! In Advances in Neural Information Processing Systems, pages 2307–2315, 2016.

[10] L. Evers and C.-M. Messow. Sparse kernel methods for high-dimensional survival data. Bioinformatics, 24(14):1632–1638, 2008.

[11] H. S. Farahani and J. Lagergren. Learning oncogenetic networks by reducing to mixed integer linear programming. PloS one, 8(6):e65773, 2013.

[12] D. Hanahan and R. A. Weinberg. Hallmarks of cancer: the next generation. cell, 144(5):646–674, 2011.

[13] F. E. Harrell, R. M. Califf, D. B. Pryor, K. L. Lee, and R. A. Rosati. Evaluating the yield of medical tests. Jama, 247(18):2543–2546, 1982.

[14] K. Jahn, J. Kuipers, and N. Beerenwinkel. Tree inference for single-cell data. Genome biology, 17(1):86, 2016.

[15] N. Karpov, S. Malikic, M. Rahman, S. C. Sahinalp, et al. A multi-labeled tree edit distance for comparing” clonal trees” of tumor progression. In 18th International Workshop on Algorithms in Bioinformatics (WABI 2018). Schloss Dagstuhl-Leibniz-Zentrum fuer Informatik, 2018.

[16] H. Kennecke, R. Yerushalmi, R. Woods, M. C. U. Cheang, D. Voduc, C. H. Speers, T. O. Nielsen, and K. Gelmon. Metastatic behavior of breast cancer subtypes. Journal of clinical oncology, 28(20):3271–3277, 2010.

[17] J. F. Lawless. Statistical models and methods for lifetime data, volume 362. John Wiley & Sons, 2011.

[18] L. Lovász. Large networks and graph limits, volume 60. American Mathematical Soc., 2012.

[19] R. G. Miller Jr. Survival analysis, volume 66. John Wiley & Sons, 2011.

[20] S. J. Pan and Q. Yang. A survey on transfer learning. IEEE Transactions on knowledge and data engineering, 22(10):1345–1359, 2009.

[21] J. Pearl. Causality: models, reasoning and inference, volume 29. Springer, 2000.

[22] D. Ramazzotti, G. Caravagna, L. Olde Loohuis, A. Graudenzi, I. Korsunsky, G. Mauri, M. Antoniotti, and B. Mishra. Capri: efficient inference of cancer progression models from cross-sectional data. Bioinformatics, 31(18):3016–3026, 2015.

[23] D. Ramazzotti, A. Graudenzi, G. Caravagna, and M. Antoniotti. Modeling cumulative biological phenomena with suppes-bayes causal networks. Evolutionary Bioinformatics, 14:1176934318785167, 2018.

[24] P. Suppes. A probabilistic theory of causality. 1973.

[25] N. Tatti. Faster way to agony. In Joint European Conference on Machine Learning and Knowledge Discovery in Databases, pages 163–178. Springer, 2014.

[26] N. Tatti. Tiers for peers: a practical algorithm for discovering hierarchy in weighted networks. Data mining and knowledge discovery, 31(3):702–738, 2017.

[27] C. H. Wagner. Simpson’s paradox in real life. The American Statistician, 36(1):46–48, 1982.

[28] A. Widodo and B.-S. Yang. Machine health prognostics using survival probability and support vector machine. Expert Systems with Applications, 38(7):8430–8437, 2011.

[29] S. Yousefi, F. Amrollahi, M. Amgad, C. Dong, J. E. Lewis, C. Song, D. A. Gutman, S. H. Halani, J. E. V. Vega, D. J. Brat, et al. Predicting clinical outcomes from large scale cancer genomic profiles with deep survival models. Scientific reports, 7(1):11707, 2017.

